# Systemic and mucosal mobilization of granulocyte subsets during lentivirus infection

**DOI:** 10.1101/2021.03.19.436235

**Authors:** Rhianna Jones, Cordelia Manickam, Daniel R. Ram, Kyle Kroll, Brady Hueber, Griffin Woolley, Spandan V. Shah, Scott Smith, Valerie Varner, R. Keith Reeves

## Abstract

Granulocytes mediate broad immunoprotection through phagocytosis, extracellular traps, release of cytotoxic granules, antibody effector functions and recruitment of other immune cells against pathogens. However, descriptions of granulocytes in HIV infection and mucosal tissues are limited. Our goal was to characterize granulocyte subsets in systemic, mucosal and lymphoid tissues during lentivirus infection using the rhesus macaque (RM) model. Mononuclear cells from jejunum, colon, cervix, vagina, lymph nodes, spleen, liver, and whole blood from naïve, and chronically SHIVsf162p3-infected RM were analyzed by microscopy and polychromatic flow cytometry. Granulocytes were identified using phenotypes designed specifically for RM: eosinophils – CD45+CD66+CD49d+; neutrophils – CD45+CD66+CD14+; and basophils – CD45+CD123+FcRε+. Nuclear visualization with DAPI staining and surface marker images by ImageStream (cytometry/microscopy) further confirmed granulocytic phenotypes. Flow cytometric data showed that all RM granulocytes expressed CD32 (FcRγII) but did not express CD16 (FcRγIII). Additionally, constitutive expression of CD64 (FcRγI) on neutrophils and FcRε on basophils, indicates the differential expression of Fc receptors on granulocyte subsets. Granulocytic subsets in naïve whole blood ranged 25.4-81.5% neutrophils, 0.59-13.3% eosinophils and 0.059-1.8% basophils. Interestingly, elevated frequencies of circulating neutrophils, colorectal neutrophils, and colorectal eosinophils were all observed in chronic lentivirus disease. Conversely, circulating basophils, jejunal eosinophils, vaginal neutrophils, and vaginal eosinophils of SHIVsf162p3-infected RM declined in frequency. Overall, our data suggest modulation of granulocytes in chronic lentivirus infection, most notably in the gastrointestinal mucosae where significant inflammation and disruption occurs in lentivirus-induced disease. Furthermore, granulocytes may migrate to inflamed tissues during infection and could serve as targets of immunotherapeutic intervention.

## Introduction

Granulocytes, which include neutrophils, eosinophils, and basophils are critical innate immune cells that form one of the first lines of immune defense against pathogens. They predominantly act against bacterial (4, 11, 19, 33, 38, 39), and parasitic pathogens (1 19, 24, 37, 39), and although granulocytes are also active against viral pathogens (2, 5, 7, 11, 14, 16, 31, 33, 37, 38), this is less well characterized. Among the subsets, neutrophils are generally the most abundant and well-characterized (4, 14, 33, 47), capable of multiple specialized methods of immune defense. In response to pathogen-induced inflammation (4, 37, 39, 47), neutrophils release cytokines, neutrophil extracellular traps (NETs), and activate phagocytosis (2, 19, 38, 39, 47). In response to viral pathogens, such as HIV, (11, 33, 37, 38), neutrophils activate NETs in the female genital tract and other tissues, perform phagocytosis and release reactive oxygen species (2, 14, 32, 38). In some cases, overly robust neutrophil responses can also cause tissue damage and further disease pathogenesis as evidenced in persons living with HIV (PLWH) and SIV-infected nonhuman primates (NHPs) (5, 14, 32, 35, 45). Conversely, granulocyte frequencies and functions may be abrogated by HIV disease (5, 13, 31), including neutrophil dysregulation and dysfunction (31) and a decline in frequency due to HIV accelerated apoptosis and hematopoietic failure in the bone marrow (5, 13, 31). Similar accelerated neutrophil apoptosis mediated by viral infection has also been observed in rhesus macaques (RM) infected with SIV (7, 8, 16).

Eosinophils and basophils can also respond to pathogens through the release of granules and cytokines, and in some cases through extracellular traps (19, 24, 26, 38, 39). While both eosinophils and basophils are primarily associated with allergens and parasitic infections, they may be effective antigen presenting cells (APCs), similar to neutrophils, and help in activating Th2 responses (21, 23, 26, 47). Additionally, eosinophils and basophils can interact with dendritic cells and B cells directly and/or indirectly through T cell activation to promote Th2 immunity, B cell proliferation and antibody production (26, 37). There has also been evidence that eosinophils play a role in limiting inflammation, tissue glucose uptake and minimizing host tissue damage (1, 37). However, similar to the effect on neutrophils, HIV may cause an expansion of eosinophils associated with a decrease in CD4+ T cells and in parasitic coinfections (31).

NHPs have long been used as models of immunological research because of their similarities to human physiology, immunology and disease pathogenesis (3, 12, 16, 20, 32). Granulocyte immune function similarities between human and NHP models include immune cell recruitment (12, 14, 24, 32), phagocytosis (16, 17, 32, 36, 39), and cytokine and chemokine release (10, 23, 32). However, phenotypic differences between human and macaque granulocytes must also be considered. In humans, CD16 is the preferred cell marker for identifying neutrophils, but NHP neutrophils do not express CD16. Thus, CD14 has previously been used to partially identify NHP neutrophils (16, 32, 47, 48). Healthy rhesus macaques can have a population of neutrophils ranging from 20-50% of all white blood cells (WBC) (12), whereas in humans this range is 40-70% neutrophils of all WBC (16, 17, 33). The frequencies of basophils (<1% of WBC) and eosinophils (0.3-10% of WBC) populations are generally similar in humans and macaques (12, 31). Regardless, consensus phenotypes for each population in NHPs, particularly in tissues, are lacking. Therefore, in this study, we performed in-depth phenotypic analysis of neutrophils, eosinophils, and basophils in systemic and mucosal tissues of naïve and SHIV-infected RM using flow cytometry and ImageStream cytometry.

## Methods

### Animal samples

Indian-origin rhesus macaques (*Macaca mulatta*) were used for all components of this study and included: experimentally naïve (n = 19), or chronically infected with SHIVsf162p3 (n = 21). Blood, biopsies and other tissues were collected where indicated and processed using standard protocols. All animals were housed at either Biomere, Inc. (Worcester, MA) or Bioqual, Inc. (Rockville, MD). All study collections were reviewed and approved by either the Biomere Institutional Animal Care and Use Committee or the Bioqual Institutional Animal Care and Use Committee.

### Tissue processing and preparation

Mucosal biopsies (colorectal and vaginal) and tissues (vagina, cervix, colon, jejunum, colorectal) were enzymatically digested with collagenase (type IV for reproductive tissues, type II for digestive tissues), pushed through 70 μm nylon mesh cell filters and then were overlaid on 35% percoll (Sigma-Aldrich) followed by 60% percoll for density gradient centrifugation. Liver tissue was mechanically disrupted and pushed through cell filters and was similarly subjected to density gradient centrifugation. The interphase containing leukocytes was collected, washed with RPMI 1640 (Corning) containing 5% fetal bovine serum (FBS), counted and used for phenotyping granulocytes. Lymph nodes and spleen were mechanically pushed through 70 μm nylon mesh cell filters followed by RBC lysis with ACK (ammonium-chloride-potassium) lysing buffer (ThermoFisher Scientific) solution. The cells were washed and resuspended in RPMI 1640 containing 10% FBS for further use in assays.

### Flow cytometric staining and analysis of whole blood and tissue samples

To identify granulocytes using polychromatic flow cytometric staining, 100μL of whole blood was stained with surface antibodies for 20 minutes at room temperature, protected from light. Antibodies against the following molecules were included: CD49d (HP2/1; Beckman Coulter), CD32 (FUN-2; BioLegend), CD95 (DX2; BD Pharmingen), CD117 (104D2; BioLegend), CD123 (6H6; BioLegend), CD20 (2H7; BioLegend), CD11b (ICRF44; BD Pharmingen), CD14 (M5E2 BD; Pharmingen), CD62L (SK11; BD Pharmingen), CD45 (D058-1283; BD Pharmingen), CD3 (SP34.2; BD Pharmingen), CD16 (3G8; BD Pharmingen), CD64 (10.1; BD Pharmingen), CD66abce (TET2; Miltenyi), CD63 (H5C6; BioLegend), HLA-DR (G46-6; BD Pharmingen), and FcRε (AER-37 (CRA-1); BioLegend). Following this, FACS lysis buffer (BD Biosciences) was used to lyse red blood cells (RBC) according to manufacturer’s recommendations. Samples were then washed twice with wash buffer (1X DPBS containing 2% FBS) and fixed with 1% formaldehyde. Cells from processed tissues were first stained with LIVE/DEAD fixable aqua stain (Life Technologies) to distinguish live cells. After 20 minutes of incubation at 4 °C, cells were washed and stained for surface markers as listed above and incubated at 4 ◻C for 20 minutes. Finally, cells were fixed using 1% formaldehyde. Data were acquired using an LSRII flow cytometer (BD Biosciences) and analyzed using FlowJo software (version 10.6.1).

### ImageStream flow cytometric staining and analysis

Whole blood and tissue mononuclear cells were stained for ImageStream cytometry similar to above with the same antibodies. Two panels were used to characterize neutrophils, eosinophils, and basophils. Markers used for the eosinophil/neutrophil panel were: CD49d, CD45, CD14, CD66abce, and HLA-DR and markers used for basophil panel were CD45, CD66abce CD123, HLA-DR, and FcRε. After 30 minutes of incubation with surface antibodies, cells were washed and stained with DAPI (Thermofisher) to visualize nuclear morphology after permeabilization with Invitrogen Fix and Perm medium kit (Thermofisher) for whole blood staining. In case of tissue immune cells, DAPI was used for live and dead cell discrimination without permeabilization. After fixation with 1% formaldehyde, samples were recorded using an ImageStreamX Mk II (EMD Millipore) and analyzed using IDEAS Application v6. The general analysis, colocalization and internalization modules in the IDEAS software were utilized in analyzing these samples.

### Statistical Analyses

Comparisons between naïve and SHIV-infected animals were analyzed by Mann-Whitney *U* tests using Prism v.8 software. Differences between test groups were considered significant when *p* < 0.05.

## Results

### Myeloid cell phenotypes in whole blood identified using flow cytometry

Initial identification of granulocyte populations (and monocytes for comparison) was determined among medium to high side scatter (granularity) CD45+ leukocytes. Subsequently cells were divided into those that express the CD66 family of receptors (eosinophils, neutrophils) and those that do not (basophils). Next, based on previous reports (16, 32, 47, 48), CD14 was used as a general marker to distinguish neutrophil populations from other subsets and each granulocyte population was defined as follows: CD45+HLA-DR-CD14+CD49d-CD66+ neutrophils, CD45+HLA-DR-CD14-CD49d+CD66+ eosinophils, and CD45+HLA-DR-CD123+CD66-basophils (Fig. 1a). Monocytes were defined as HLA-DR+CD14+CD16+/− to provide discrimination of all myeloid cell phenotypes in whole blood of RM (Fig. 1 and 1b). To further confirm phenotypic identity in RM, Fc receptor expression was compared among all populations. As expected, all subsets had high frequencies of CD32 (FcRγII; Fig.1b). CD64 (FcRγI) was expressed at variable levels on all populations except basophils, but as expected, basophils were the only cells to demonstrate high constitutive expression of FcRε. In strong agreement with previous studies for RM (18), CD16 (FcRγ III) was not expressed on granulocytes but only on a subset of monocytes. Cell frequencies in whole blood of experimentally naïve RM were analyzed to evaluate baseline frequencies (Fig. 1c), and as expected, neutrophils had the highest frequency out of the three subsets (median 51.1, range 25.4-81.5%), followed by eosinophils (median 3.16, range 0.59-13.3%) and basophils (median 0.048, range 0.059-1.8%).

**FIG1:**
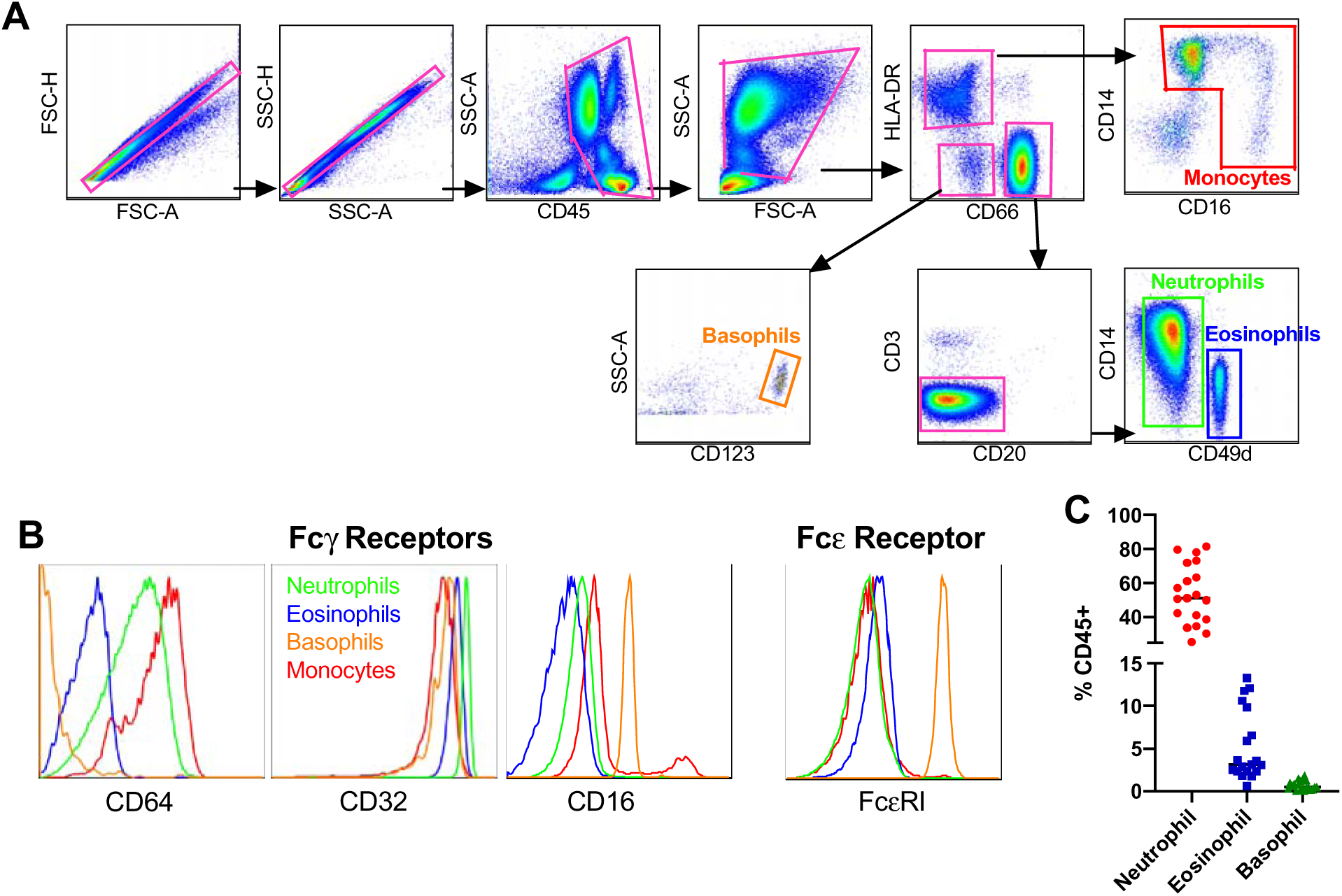
Flow cytometric analysis of whole blood granulocyte surface markers. Flow cytometric panel determining granulocyte subpopulations (A) shown using whole blood samples. Frequencies of CD64, CD32, CD16, and FcRε receptors on neutrophils (green), eosinophils (blue), basophils (yellow), and monocytes (red), in naïve rhesus whole blood samples (B). Baseline frequencies of granulocyte subsets in rhesus whole blood samples (C): neutrophils (red), eosinophils (blue), and basophils (green) determined by flow cytometry.

### Granulocyte morphology and surface phenotypes by ImageStream cytometry

To further confirm the identity of RM granulocyte subpopulations, we next evaluated bulk and individual cells phenotypically and morphologically by ImageStream cytometry. Whole blood samples were stained for DAPI to identify nuclear morphology and surface markers to confirm surface expression of markers used in the standard flow cytometry panels. Using the general analysis wizard of the IDEAs Application v6, DAPI+ cells were selected and representative analyses are shown in Figure 2. As in Figure 1, neutrophils were confirmed by surface co-expression of CD14 and CD66, eosinophils by co-expression of CD49d and CD66, and basophils by co-expression of FcRε and CD123 (Fig. 3). Further analysis of internalization in the IDEAs software confirmed the nuclear morphology of the granulocyte populations. Analysis of DAPI+ cells showed multilobed nuclei of CD14+CD49d-CD66+ neutrophil populations, bilobed nuclei of CD14-CD49d+CD66+ eosinophils and granular cytoplasm of basophils as expected (Fig. 3).

**FIG 2:**
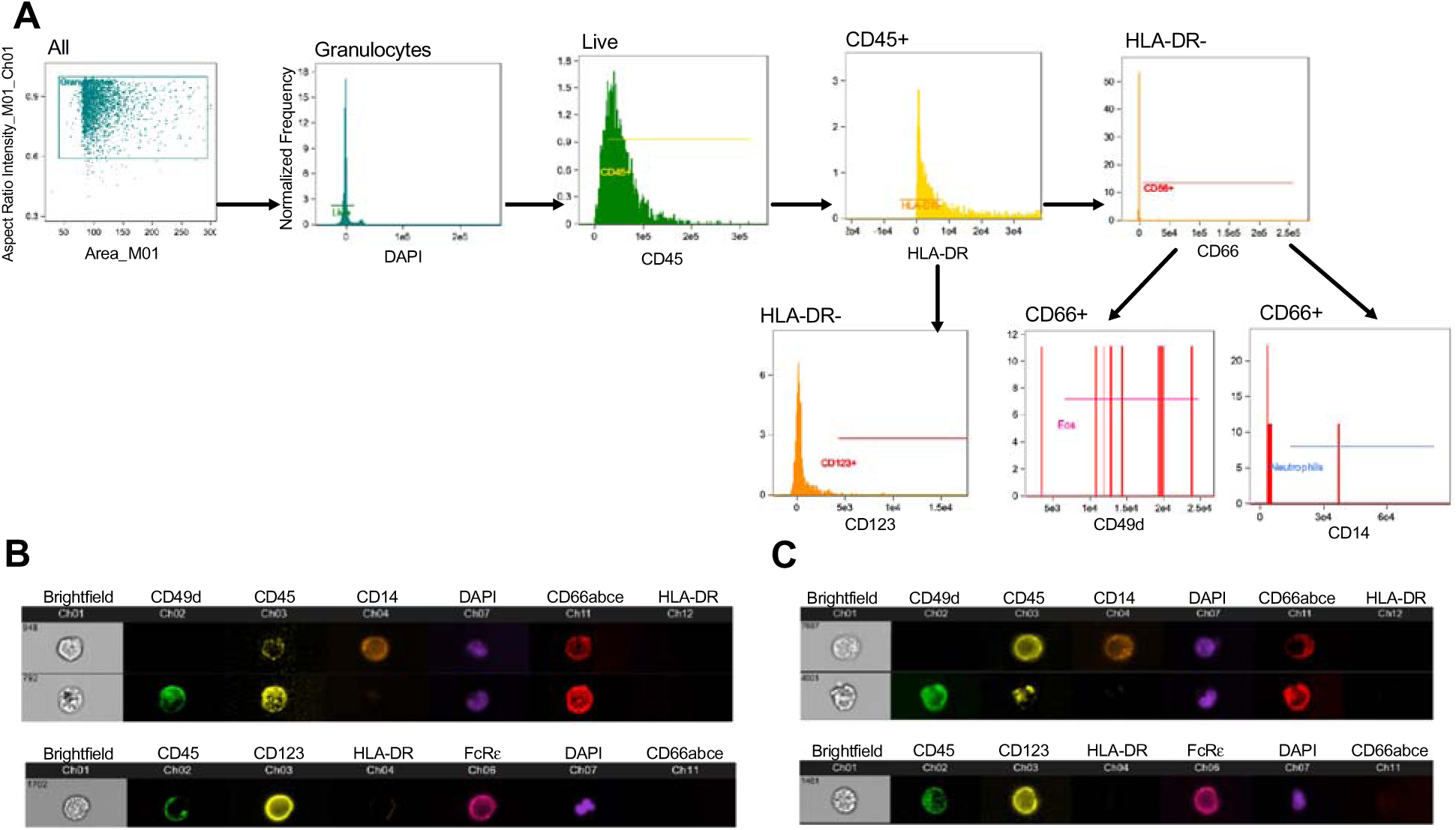
Granulocyte phenotypes analyzed by ImageStream. ImageStream flow cytometry gating strategy (A) used to determine populations of neutrophils, eosinophils, and basophils in whole blood. Image representation of (B) neutrophils (top), eosinophils (middle), and basophils (bottom) isolated from rhesus whole blood.

**FIG 3:**
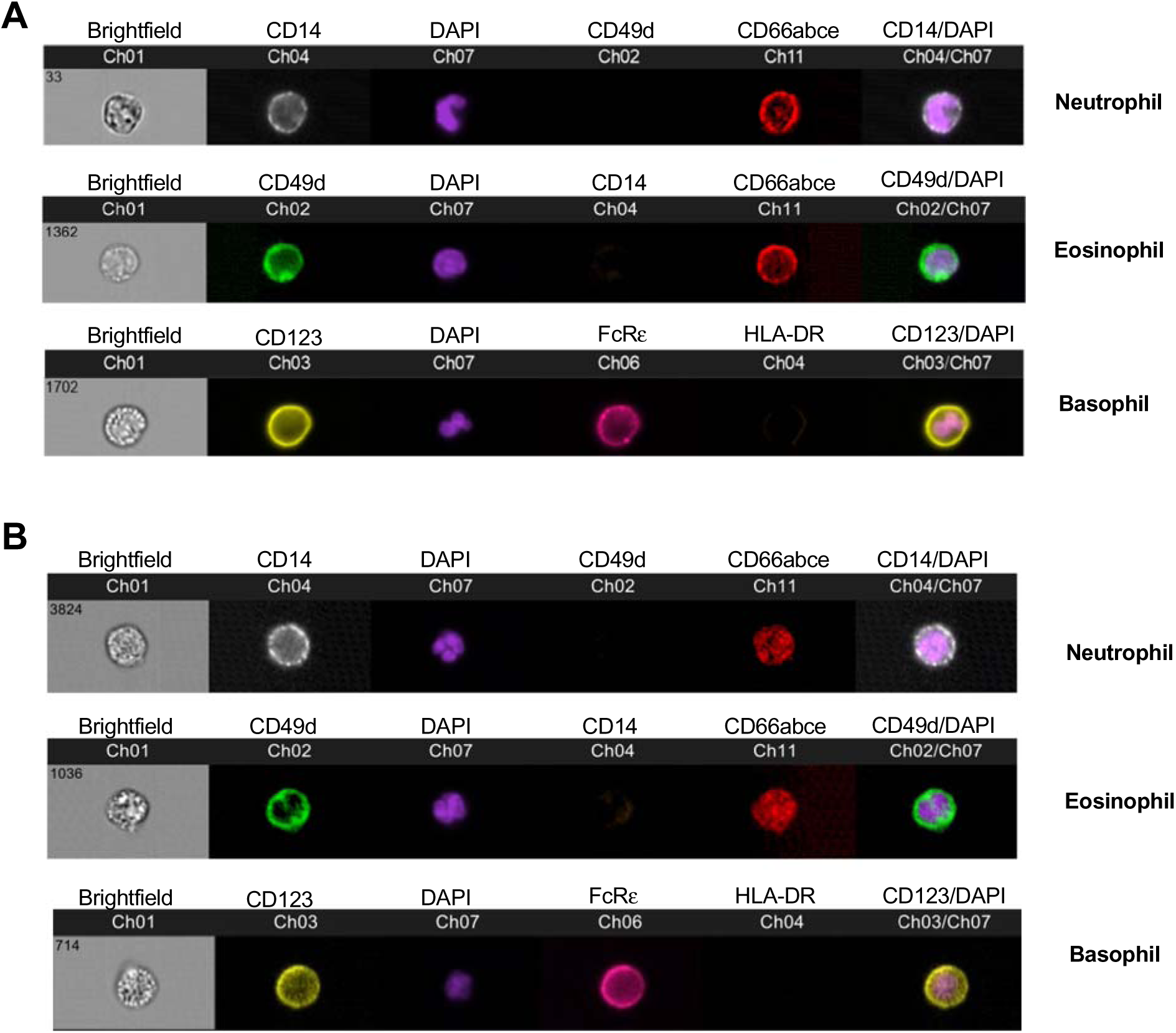
Nuclear morphology of granulocyte subsets analyzed by ImageStream. ImageStream flow cytometry images generated from whole blood samples and two representative evaluations are shown (A and B). Internalized DAPI for visualizing nuclear morphology is overlaid with surface markers indicating neutrophils (top), eosinophils (middle), and basophils (bottom).

### Modulation of peripheral and tissue granulocytic subsets in infected animals

In a cross-sectional evaluation of naïve RM and chronically SHIV-infected RM, we first quantified granulocytes in whole blood. Circulating neutrophils were increased over 50% in SHIV-infected animals compared to naïve RM (medians, 79.6% versus 51.1%). While circulating eosinophils were largely unaltered, basophils were modestly, but significantly (p = 0.003) reduced in infected animals (Fig 4).

**FIG 4:**
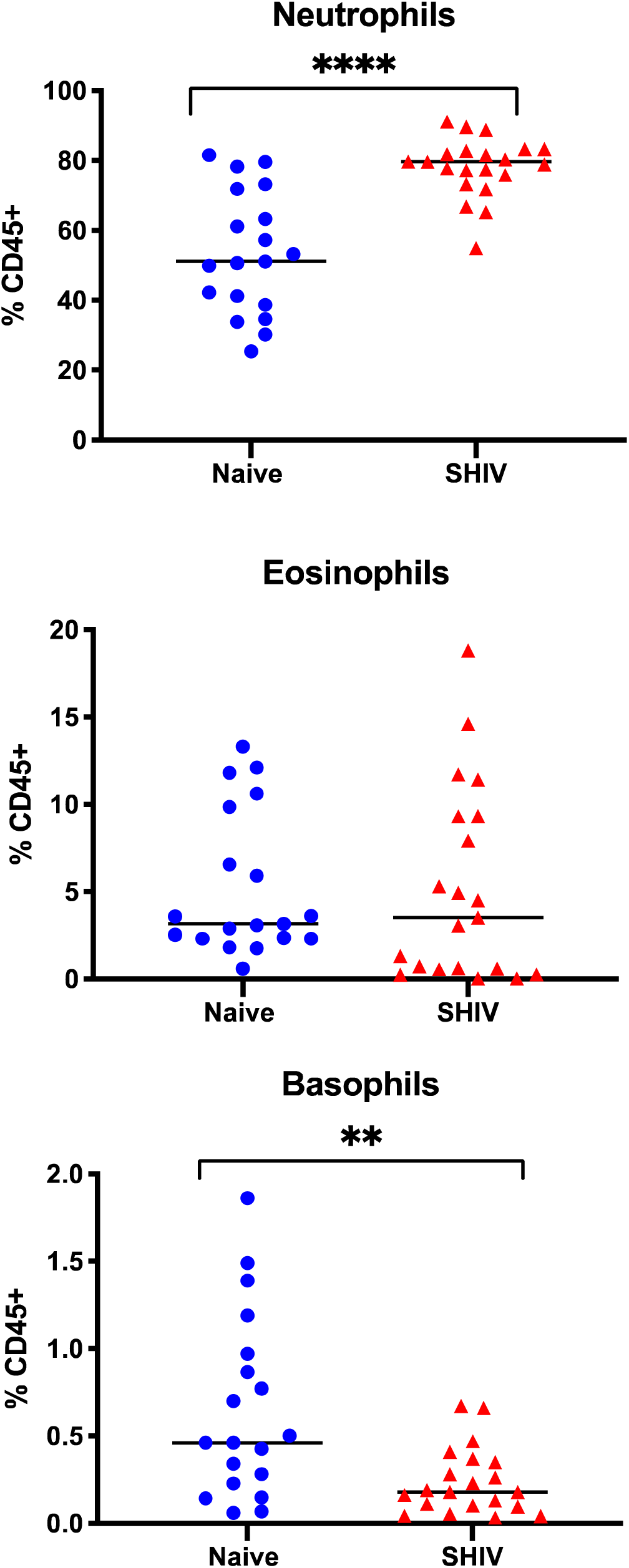
Granulocyte subsets in whole blood of naïve and lentivirus-infected rhesus macaques. Granulocyte subset frequencies determined in peripheral blood by flow cytometry. Chronic SHIV infection whole blood samples (red) are compared to naïve (blue). ** P< 0.01, **** P <0.0001, Mann-Whitney test.

Next we applied our analyses of granulocytes in whole blood to tissues collected by mucosal biopsy or at necropsy from both naïve and chronically SHIV-infected animals and observed neutrophils and eosinophils in vaginal biopsies of naïve animals (medians, 0.65% and 0.38% respectively) (Fig. 5a). Interestingly, in SHIV-infected RM, both cell types were significantly reduced. Neutrophils and eosinophils were both found at low frequencies in colorectal biopsies of naive animals (medians, 0.083% and 0.021% respectively), but were significantly increased during infection (both 3-fold increases).

**FIG 5:**
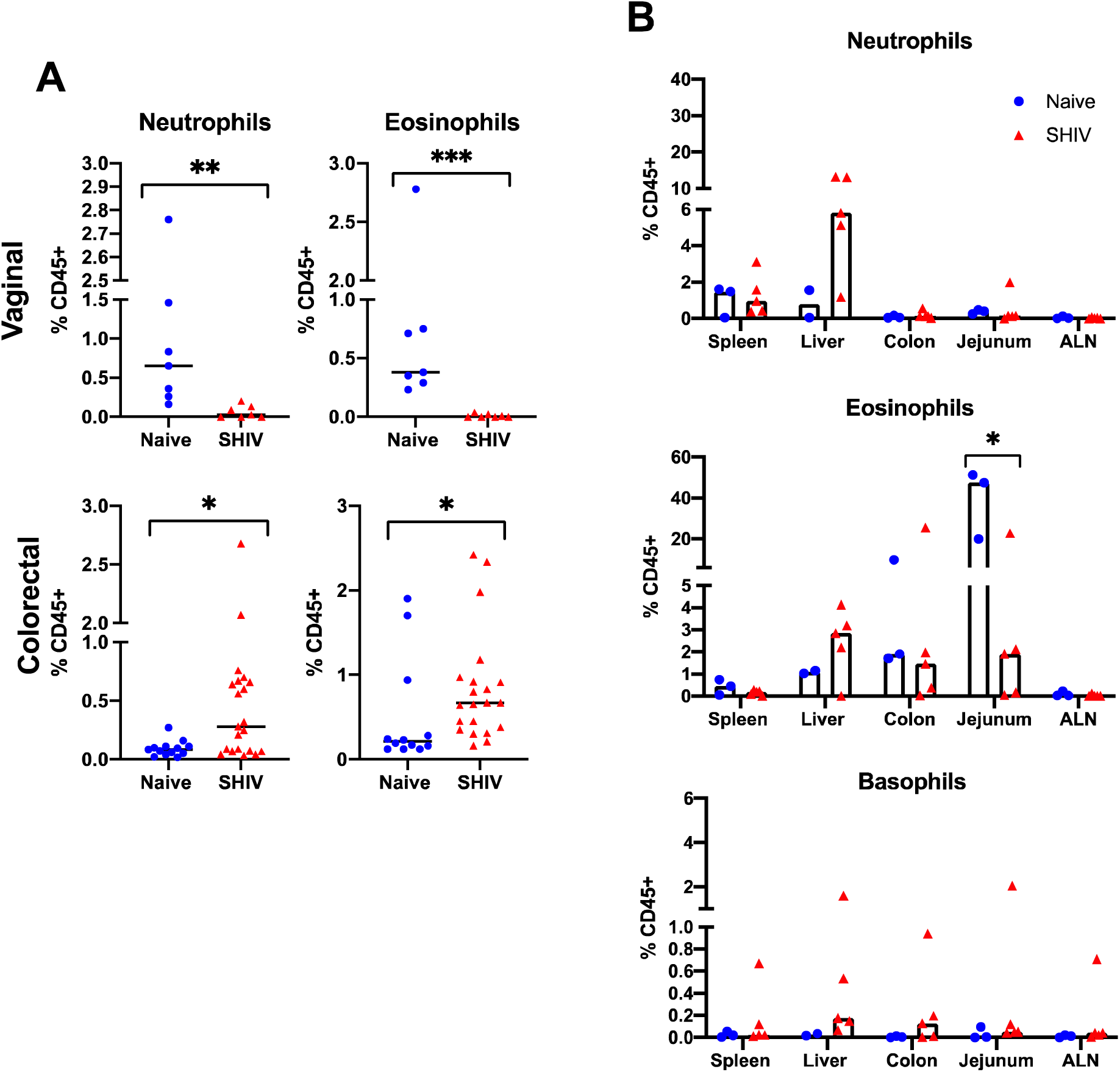
Granulocyte subsets in mucosal and lymphoid tissues of naïve and lentivirus infected macaques. Frequencies (A) of neutrophils (top), eosinophils (middle), and basophils (bottom) found in naïve (blue) and chronic SHIV (red) infected rhesus macaque tissues. Frequencies (B) of neutrophils (left) and eosinophils (right) found in vaginal (top) and colorectal (bottom) biopsies from naïve (blue) and chronic SHIV (red) infected rhesus macaques analyzed using flow cytometry. * P< 0.05, ** P< 0.01, *** P< 0.001, Mann-Whitney test.

Quantification of granulocytes in a very small subset of the animals in spleen, liver, colon, jejunum, and axillary lymph nodes from necropsy tissues revealed detectable populations in most animals (Fig. 5b). Following infection we were unable to detect significant changes, most likely due to the low number of animal samples. In particular, basophil frequencies were near the threshold of detection in tissues, and aside from one animal these frequencies were also unchanged during infection. Notably, both eosinophils and neutrophils seem to increase in liver tissue, but larger studies will be needed to validate this finding. We also noted an interestingly high frequency of eosinophils in jejunum of naive animals (median, 47%) which could also be of interest in future studies.

## Discussion

Granulocytes play major roles as innate immune cells against bacterial, viral and parasitic pathogens which have been described in multiple studies (11, 19, 24, 33, 38, 39). Their protective role against HIV as well as their pathogenic role causing tissue damage in HIV infected patients, particularly by neutrophils, have been reported in multiple studies (12, 16, 17, 31, 33, 47, 48). However, NHP granulocytes in mucosal tissues of chronic SHIV infections have been underexplored. Here, we describe granulocyte subset modulations in systemic and mucosal tissues of SHIV infected RM.

Using flow cytometry, we first identified the phenotype of granulocytes as CD14+CD49d-CD66+ neutrophils, CD14-CD49d+CD66+ eosinophils, and CD123+CD66-basophils. A major difference in phenotype between human and NHP neutrophils is the expression of CD14 by NHP neutrophils rather than CD16 as seen in human neutrophils. Indeed, a subset of monocytes were the only myeloid cells to express CD16 in NHP. Due to the limited availability of cross-reactive antibodies to further characterize granulocytic subsets, we confirmed the granulocyte phenotype by examination of their nuclear morphology and surface marker expression. ImageStream, a tool that combines microscopy and flow cytometry, allowed us to visualize the polymorphonuclear neutrophils, bilobed eosinophil nuclei and granular basophils (Fig. 2 and 3), thus confirming the granulocyte phenotypes we have identified for rhesus macaques.

Persistent immune activation is commonly associated with chronic HIV infection. Studies examining neutrophils in both acute and chronic SIV infection (13, 32) showed altered functional and phenotypic differences in CD11b+ neutrophils, however a reduction of these cells were seen in acute infection. Similarly, in people living with HIV, neutropenia (22, 29, 30, 41, 46), and impaired phagocytic activity and cytokine production (10, 27, 30, 36) have been frequently observed. Interestingly, we observed significantly increased neutrophils and decreased basophils compared to naïve animals in the blood of SHIV-infected animals. While basophils are predominantly located in the blood, our observed decline in basophils may be due to their infiltration of inflamed tissues as seen in the gastric mucosa of *Helicobater pylori* infected patients (6) and in the skin of the cutaneous allergic reactions (40).

Progressive HIV infection can cause immune dysfunction in mucosal tissues of the gut and also the reproductive tract (14, 15, 43, 44, 50). In this study we found elevated frequencies of neutrophils and eosinophils in colorectal biopsies and higher eosinophil frequency in jejunum samples of chronic SHIV infected animals compared to naïve animals (Fig. 5) indicating granulocytic infiltration of gut tissues. Similar observations of increased neutrophil infiltration and epithelial barrier dysfunction have been reported in the gut tissues of SIV infected animals and treated and untreated chronic HIV infected patients (9, 45). Further, Hensley-McBain et al have showed that HIV induced microbial translocation and dysbiosis characterized by reduced *Lactobacillus: Prevotella* ratio as the reason for the increased neutrophil recruitment and increased neutrophil lifespan in the colorectal mucosa of HIV infected individuals (15). Neutrophil infiltration in the colonic mucosa is also correlated with disease severity in inflammatory bowel disease, thus further suggesting that elevated neutrophils in the gut are potentially inflammatory and induce additional mucosal damage.

Interestingly, chronic SHIV infected vaginal biopsies had significantly lower frequencies of neutrophils and eosinophils than naïve animals in our study. This is in accordance with studies that have shown increased neutrophil apoptosis in whole blood and tissues, and eosinophil apoptosis in whole blood during lentivirus infection in humans (7, 8, 16, 45, 49). Similarly in PLWH vaginal inflammation was associated with increased neutrophils (33). *In vitro* data suggests that neutrophils are capable of entrapping and inactivating HIV through NETS within minutes of viral exposure (2). Based on this, our data suggests differential modulation of granulocyte trafficking, expansion and functions of resident granulocyte subsets in different mucosal tissues by chronic SHIV infection. However, more research exploring granulocyte functions in mucosal tissues, particularly in chronic SIV/SHIV infection is warranted.

In summary, our study provides comprehensive data on granulocyte phenotypes in blood and tissues of RM, which was consistent with other reports (16, 25, 32). Visualization of granulocytes by ImageStream proved to be an excellent tool to confirm immune cell phenotypes. Further, our data showed that mucosal tissues which are the sites of viral infection show modulated frequencies of granulocyte subsets and thereby could differ in their functional outcomes of either immunoprotection or tissue damage. While we analyzed other systemic and lymphoid tissues including spleen, liver and axillary LN, we did not see any changes. The limitations of our study include a small sample size and availability of only cross-sectional data for chronic SHIV infection. Longitudinal time points in chronic SHIV infected animals would provide a better understanding of the granulocyte subset modulations. Overall, our data provides evidence of immune modulation of granulocytes in gut and vaginal tissue and warrants more research into identifying their roles in protection/ pathology in lentivirus infection.

## Acknowledgements

The efforts of the authors were supported by NIH grants R01 DE026327, P01 AI120756, R21 AI145678 (all to R.K.R.), as well as the Harvard University Center for AIDS Research Advanced Laboratory Technologies Core (P30 AI060354). The authors also wish to thank Dr. Dan Barouch for providing some animal samples and Michelle Lifton for assistance with flow cytometry.

## Abbreviations

RM: Rhesus macaques
HIV: Human Immunodeficiency Virus
SIV: Simian Immunodeficiency Virus
SHIV: Simian-Human Immunodeficiency Virus

